# Sintering 3D-Printed Hydroxyapatite-Wollastonite Lattices Improve Bioactivity and Mechanical Integrity for Bone Composite Scaffolds

**DOI:** 10.1101/2025.04.06.647463

**Authors:** Peter M. Bertone, Levi M. Olevsky, Kavin Kathir, Simon A. Agnew, William J. Scheideler, Katherine R. Hixon

## Abstract

The advancement of bone tissue engineering relies on the development of scaffolds that combine structural integrity with bioactivity. This study introduces a novel composite scaffold integrating three-dimensional (3D) printed hydroxyapatite (HA)-wollastonite (WOL) gyroid lattices with chitosan-gelatin cryogels, designed to fulfill these dual requirements. The HA-WOL lattices were fabricated using digital light processing (DLP) 3D-printing and subjected to optimized thermal treatment cycles demonstrating statistically superior compressive modulus and ultimate strength. This thermal process facilitated the phase transformation of HA-WOL to bioactive β-tricalcium phosphate (β-TCP) and silicocarnotite mixed phases, with MG63 (osteoblast-like) cell culture revealing significantly enhanced viability and biocompatibility. The chitosan-gelatin polymer network was successfully incorporated into the lattice, resulting in a composite scaffold with retained relative swelling capacity, improved mechanical stability, and superior bioactivity compared to cryogel-only constructs. Additional MG63 cell culture studies revealed that the composite scaffold supported cell viability and proliferation into the constructs, demonstrating its potential to conduct tissue regeneration across bone defects. This work highlights the synergistic effects of integrating bioactive ceramics with polymer-based cryogels, offering a promising solution to address bone regeneration in orthopaedic reconstruction. Future research will focus on *in vivo* validation and optimization of scaffold architecture to further enhance clinical relevance. This study paves the way for next-generation composite scaffolds capable of bridging the gap between mechanical integrity and biological performance in bone regeneration.

## 1. Introduction

Bone is the second most transplanted tissue in the United States, with bone defects most often arising from acute trauma, tumor resection, and/or infection, necessitating treatment with bone grafts.^1,2^ Further, with both an increasing life expectancy and incidence of bone grafting procedures, there is a growing focus on improving orthopaedic interventions and their outcomes.^3,4^ Bone is a complex tissue composed of both organic and inorganic materials. The organic matrix, primarily consisting of collagen fibers, forms a porous framework that provides flexibility and tensile strength. The inorganic mineral phase, predominantly hydroxyapatite (HA), imparts hardness and load-bearing capacity. Together, these components enable bone to perform its crucial structural and metabolic functions.^2,5^ Given bone’s complex structural and regional heterogeneity, autologous bone transplantation remains the gold standard for bone grafting; however, this approach has significant limitations including donor site morbidity, pain, limited tissue availability, multiple surgeries, and an increased risk of infection.^2,6^ Furthermore, these challenges are exacerbated in complex anatomical sites, such as long bones or the maxillofacial region, underscoring the need for alternative approaches to regenerate bone tissue effectively.^7,8^

Recent advancements in biomaterials have significantly accelerated the development of alternative bone graft substitutes.^2,9,10^ Ideally, a bone-targeted biomaterial should create a conducive environment for endogenous cells to lay down new bone while degrading at a rate compatible with this tissue formation, simultaneously emulating the morphology, porosity, and load-bearing capacity of healthy, native bone tissue. While many tissue-engineered materials possess some combination of these requisite characteristics, cryogel scaffolds are a notable standout.^11^ Cryogels are polymeric biomaterials fabricated via a tailorable freeze-thaw cycle at sub-zero temperatures to produce a sponge-like, macroporous structure that is non-toxic, fully resorbable, and mimics cancellous bone morphology.^11,12^ In particular, our group and others have worked extensively with fabricating and optimizing naturally derived polymer cryogels made from chitosan (Ch) and gelatin (Gn) for clinical use as a bone graft substitute.^13–15^ However, while mechanically durable and resilient, cryogel utility as a bone graft substitute is hindered by its low ultimate strength.^16^ Further, cryogels alone are not sufficiently stiff nor stable enough to integrate directly with the metallic implants often required for defect and bone graft fixation. Comparatively, 3D-printing has been used as a viable method for bone tissue engineering scaffold fabrication.^17,18,19^ While there are several techniques for 3D-printing, our group specifically utilizes digital light processing (DLP) printing to fabricate scaffolds for bone. DLP printing is particularly advantageous in targeting bone tissue scaffolds due to its ability to produce high-resolution structures with precise control over shape, pore size, and interconnectivity, enabling fabrication of structures that can be patient-specific and integrate with clinical hardware.^20,21^ These characteristics are crucial for creating customized lattices that mimic the complexity of natural bone architecture while working effectively in a clinical context. Previous work from our group demonstrated successful cryogelation of Ch-Gn within plastic 3D-printed frameworks (gyroid lattice).^22^ The resultant combination scaffolds were measured to be over 1000 times stronger than cryogel alone via ultimate compression. While this work demonstrated successful fabrication with a significant increase in mechanical stiffness, it was observed that the proprietary FormLabs 3D-printed structures had a long-term negative effect on cells, with complete cell death by day 14 due to cytotoxic organic components within the resin.^22^ In addition to improving the 3D-printed structure biocompatibility, an ideal alternative resin material should also provide distinct benefits including increased scaffold bioactivity and mineralization (e.g., calcium-phosphate and calcium-silicate minerals).

Hydroxyapatite (HA; Ca_10_(PO_4_)_6_(OH)_2_) and wollastonite (WOL; CaSiO_3_) are naturally occurring calcium-phosphate and calcium-silicate minerals, respectively. HA serves as the primary inorganic component of bone and teeth, while WOL is naturally found in rock deposits or can be synthetically produced.^23–25^ Categorized as a bioactive and nontoxic ceramic, HA has been extensively utilized in bone tissue engineering due to its ability to induce osteogenic differentiation and promote osteoblast proliferation.^26^ Synthetic HA materials, with chemical and structural properties analogous to native bone, have been demonstrated to enhance the adhesion and proliferation of osteoblast cells on their surfaces.^27^ However, depending on its density, HA is characterized by poor resorption rates which can limit its effectiveness in biomedical applications, as materials that fail to resorb can impede tissue remodeling and integration. WOL, on the other hand, has been shown to improve the mechanical, bioactive, and degradation properties of biomaterials when combined with HA.^28,29^ Consequently, directly combining cryogel with HA-WOL ceramic lattice has the potential to create polymer macroporous composites with enhanced bioactivity and regenerative capabilities for bone tissue engineering. While cryogels have previously been formed with mineral particles^27,30–32^ and integrated with a polymer 3D-printed PTMC-HA gyroid,^33^ no studies have formed the composite scaffolds with only inorganic mineral materials. In such a composite, the cryogel would offer a highly porous structure supporting cell ingrowth and proliferation, while the ceramic lattice component would provide robust structural support while promoting both osteo-conduction, and -induction, facilitating effective bone tissue formation. Despite these advantages, HA printed structures are inherently brittle^34^ and lack desired resorption properties, while DLP resins contain cytotoxic organic components which restrict their use in medical applications.^35^

Decomposition and sintering are well-known thermal processes used to modify the physical and bioactive properties of ceramics,^36^ including HA and WOL.^37–39^ Researchers have shown that initial decomposition at moderate temperatures will burn off organic binders and volatile substances present in the resin, leaving behind a body composed primarily of constituent ceramic particles.^40^ This is followed by a sintering process at higher temperatures, which facilitates the densification and consolidation of the ceramic particles, enhancing the mechanical strength and structural integrity of the scaffold.^39,41^ Controlled sintering parameters are not only useful in forming more homogenous ceramic composites but have also been shown to improve the material’s bioactivity, as well as osteo-conduction and -induction properties.^42^ Thus, these thermal treatments are pivotal in transforming 3D-printed ceramics into robust, bioactive structures that closely mimic the inorganic components of bone, offering significant potential for advanced bone tissue engineering applications.

While cryogel, HA, and WOL scaffolds have been studied separately, the integration of 3D-printed HA-WOL lattice with cryogel remains unexplored.^39^ This study aims to investigate the effects of sintering on the physical and bioactive properties of DLP-printed HA-WOL structures and evaluate the efficacy of combining Ch-Gn cryogels with 3D-printed HA-WOL lattices. We aim to determine if this integrated approach can overcome the limitations of each material when used individually, yielding a composite with superior mechanical properties and enhanced biological performance. Therefore, we hypothesize that sintering DLP-printed HA-WOL lattices will significantly enhance mechanical strength and bioactivity while reducing cytotoxicity compared to green lattices (printed and cured). To test this, we quantified the effects of two sintering cycles on material composition, crystallinity, and mechanical strength of the printed HA lattices, and evaluated their bioactivity by assessing osteoconductive properties with MG63 (osteoblast-like) cells. Finally, we propose that integrating sintered HA structures with Ch-Gn cryogels will further improve the scaffold’s physical and mechanical properties, bioactivity, and osteoconductive potential when compared to individual components (i.e., HA-WOL lattice or Ch-Gn cryogels alone). To explore this, we characterized the composite scaffold’s porosity, morphology, mineralization potential, and mechanical characteristics, alongside cell proliferation and infiltration in a 3D culture over 14 days. Overall, this work provides a novel approach to designing, fabricating and treating advanced biomaterials that integrate bioactivity, resorbability, mechanical strength, and biocompatibility, addressing key limitations of current bone graft substitutes. By combining sintered HA-WOL lattices with cryogel, this work aims to lay the foundation for exploring more effective bone tissue engineering strategies, with the potential to improve clinical outcomes.

## 2. Materials and Methods

### 2.1 Fabrication of 3D-printed lattices

3D ceramic lattices and bulk material cylinders were printed on a Bison 1000 DLP Printer (Tethon 3D, Omaha, NE, USA) using Osteolite resin (Tethon 3D). The printing process involved a basic exposure time of 38 seconds, an initial exposure time of 50 seconds, and a light engine brightness set to 200 to achieve 100 μm layer thickness. After printing was complete, the pieces were detached from the build platform with a smooth-edged spatula. The supports were then manually removed, and the pieces cleaned in an ultrasonic cleaner (Branson 200, Branson Ultrasonics, Brookfield, CT, USA) using isopropyl alcohol to eliminate any remaining non-polymerized suspension. Once dry, the pieces were exposed to an ultraviolet (UV) lamp for 15 minutes to cure.

### 2.2 Thermal treatment of 3D-printed lattices

Thermogravimetric Analysis (TGA) was performed on the lattices to determine the temperature at which the resin binder was predominantly burned away (Supplemental Fig. 1). This temperature was found to be around 465°C (TGA 550, TA Instruments, New Castle, DE, USA). Then, two different de and sinter cycles were tested:

- <u>Cycle 1</u>: 0.83°C/min until 232°C, hold 8 hours, 1.25°C/min until 700°C, hold 1 hour, then cool to room temperature in table top furnace (ThermoLyne, Waltham, MA, USA) and transfer to muffle furnace (1400GCF Muffle Furnace, Across International, Livingston, NJ) for 1.66 °C/min until 1050°C, hold 1 hour, cool 5°C/min for 3.6 hours.
- <u>Cycle 2</u>: 1 °C/min from 50°C to 600 °C, hold 1 hour, Cool 5°C/min in tabletop furnace and transfer to muffle furnace for 10°C/min to 1250°C, hold 5 hours, cool 5°C/min to room temperature.^43^

### 2.3 Fabrication of Cryogels and Composite Scaffolds

#### 2.3.1 Formation of Cryogels

Ch-Gn cryogels were prepared based on previously established methods [10]. Briefly, a 10 mL aliquot of 1% acetic acid (Fisher Scientific, Fair Lawn, NJ, USA) in deionized water (DI) was divided into 8- and 2-mL portions in scintillation vials. Low-viscosity chitosan (80 mg, Mw = 1526.464 g/mol, MP Biomedicals, Solon, OH, USA) was added to the 8 mL portion and vortexed for 30 seconds, followed by mechanical spinning for 1 hour. Gelatin from cold water fish skin (320 mg, Mw = 60 kDa, Sigma-Aldrich, St. Louis, MO, USA) was then added to the 8 mL portion and mechanically spun for 1 hour to ensure complete dissolution. The remaining 2 mL of 1% acetic acid was mixed with glutaraldehyde (Sigma-Aldrich, St. Louis, MO, USA) to form a 1% glutaraldehyde solution. Both the 8 mL and 2 mL vials were then refrigerated at 4°C for 1 hour. Pre-cooled 3-mL syringes were filled and immediately placed in an ethanol bath within a −23°C freezer for 18 hours to crosslink at subzero temperatures.

#### 2.3.2 Formation of Cryogels within 3D-printed lattices

Syringe barrels (3 mL, Fisher Scientific, Fair Lawn, NJ, USA), containing two sintered ceramic lattices each, were pre-frozen at −23°C for 18 hours. The previously mentioned 8- and 2-mL solutions were then combined by decanting between the vials and immediately poured into the pre-cooled syringes. After filling the syringes, the plungers were inserted, the syringes were flipped upside down, and the plunger was slowly pressed until the cryogel solution fully infiltrated the gyroids, allowing some excess solution to be expelled if necessary. Immediately afterward, all syringes were frozen the same as cryogels. Cryogels and the lattice containing cryogel (composite scaffolds) were lyophilized (FreeZone Freeze Dryer, Labconco, Kansas City, MO, USA) for 24 hours and stored in a desiccator until use (Fisherbrand Acrylic Desiccator, Waltham, MA, USA).

### 2.4. Characterization of lattices and scaffolds

#### 2.4.1 Morphology and porosity

Scanning Electron Microscopy (SEM, VEGA3, TESCAN, Brno, Czech Republic) was used to both visualize material morphology for all samples, and measure porosity of the cryogelated samples. Image analysis software, PoreVision, was used to measure and quantify sample average pore size.^44^ Energy Dispersion Spectroscopy (EDS/EDX, Octane Elect, EDAX, New Jersey, USA) was used to measure elemental density in each sample. Samples undergoing SEM and EDS were mounted on aluminum stubs and sputter-coated with gold for 240 seconds at 15 mA using a HUMMER 6.2 (Anatech, Sparks, NV, USA).

#### 2.4.2 Solvent dissolution

Solvent dissolution was performed to quantify the mineral content within the resin. A pre-weighed sample of resin was dissolved in 100% isopropyl alcohol and then agitated via a vortex shaker until the solution was homogenous. Once fully dissolved, the solution was filtered through a fine filter (∼10-20 μm) to isolate the undissolved mineral fraction. The residue was then rinsed with additional isopropyl alcohol solvent to remove any residual resin. The mineral fraction was air-dried at room temperature and subsequently weighed to determine the mineral content as a percentage of the initial resin sample.

#### 2.4.3 Chemical and crystalline composition

Fourier-transform infrared spectroscopy (FTIR) and X-ray diffraction (XRD) were used to compare sintered and green lattice chemical and crystalline composition, respectively. Printed samples were crushed into a fine powder using a mortar and pestle for analysis. FTIR was used (FT/IR-6200, Jasco, Oklahoma City, OK, USA) with an attenuated total reflectance (ATR) accessory to analyze the chemical composition of printed pieces. Powder XRD was performed using a Rigaku MiniFlex diffractometer (Rigaku Corporation, Tokyo, Japan) with Cu Kα radiation and a scanning rate of 5° per minute. Spherical grain size estimates were calculated using the Debye-Scherrer method,

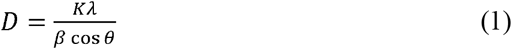

where *D* is crystallite size, *K* is the shape factor (assuming 0.98), λ is the X-ray wavelength (Cu Kα, 1.54 Å), β is the FWHM of the diffraction peak in radians, and θ is the Bragg angle.^45^

#### 2.4.4 Mechanics

Mechanical testing was performed using an Instron 68SC-2 system (Instron, Norwood, MA, USA) with a 500 N load cell and a test rate of 10 mm/min, preload of 0.05 N, and preload speed of 1mm/min. Data was collected on Bluehill universal software (Instron), and the compressive modulus (MPa) was calculated using Microsoft Excel.

#### 2.4.5 Swell kinetics

Swell testing was performed to evaluate the shape retention and rehydration potential of the scaffolds. Lyophilized for 24 hours prior to recording the dry weight, each sample was then placed in 5 mL of DI water and weighed at nine time points: 2, 4, 10, 20, 40 min, 1, 2, 4, and 24 h. The absolute swelling ratio and the relative swelling ratio were calculated using the following equations:

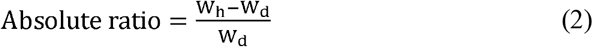

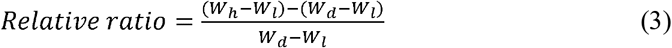

where W_h_ is the hydrated weight, W_d_ is the dry weight, and W_l_ is the weight of the lattice.^13^ The weight of the lattice was then subtracted from the final weights to isolate the swelling that is derived from only the cryogel portion of the composite scaffold.

#### 2.4.6 Mineral formation

To assess mineral induction capability, lattices and scaffolds samples were soaked in 200 mM of CaCl_2_ for 20 minutes, rinsed in DI water, soaked in 120 mM of Na_2_HPO_4_ for 20 minutes, and rinsed (repeated five times).

### 2.5 Material biocompatibility and cell bioactivity

Osteoblast-like cells (MG63, passage 94, ATCC, Manassas, VA, USA) were used for all *in vitro* cell studies. Lattices, cryogels and composite scaffolds were sterilized in 70% ethanol (Fisher Scientific, Fair Lawn, NJ, USA) for 30 min, followed by three, ten-min washes with sterile phosphate buffered saline (PBS). Cells were cultured in complete media (Dulbecco’s Modified Eagle’s Medium (DMEM, ThermoFischer Scientific, Waltham, MA, USA), 10% fetal bovine serum (Omega Scientific Inc., Tarzana, CA, USA), and 1% penicillin-streptomycin solution (Life Technologies Corporation, Carlsbad, CA, USA)) until confluency was reached.

#### 2.5.1 Live/Dead stain

Following sterilization, lattices were placed in sterile, non-treated 24-well plates (Falcon, Marlboro, NY, USA), and their surface was seeded dropwise with 10,000 cells in 100 µL and left to incubate for one hour at 37°C and 5% CO_2_ to ensure cellular attachment. Then, an additional 200-400 μL of complete media was added to ensure the lattices were submerged. A treated 24-well plate was seeded with the same number of cells and evaluated at the same time points, serving as a control. Cell viability and early cell adhesion was assessed with a Live/Dead cell kit (AAT BioQuest, Pleasanton, CA, USA) After 6, 24, and 48 hours of co-culture, 500 μL of combination dye was added to the wells containing lattices and controls and stained for 30 minutes in the incubator. Cells were detected using a fluorescence microscope (EVOS M5000 Imaging System, Waltham, MA, USA), and images were taken of both the wells containing the lattice, and of lattice-only following removal from the seeding well. Live cells appeared green, whereas compromised, dead cells appeared red.

#### 2.5.2 Cytotoxicity

Cytotoxicity was measured using a Cell-Counting-Kit-8 (CCK-8, Sigma-Aldrich, St. Louis, MO, USA). Lattice cell seeding was the same as described previously. After 6, 24, and 48 hours, 300 μL of DMEM containing 10% CCK-8 solution was added to each well. After 2-3 hours of incubation, 100 μL of the solution was taken from each well and transferred to a 96-well plate from which absorbance was measured at 450 nm (n=3/group/timepoint) with a microplate reader (SpectraMax, San Jose, CA, USA).

#### 2.5.3 Cell adhesion, morphology, and proliferation

Following the same lattice seeding technique and culture procedure as above, samples were seeded with a density of 50,000 cells/100 μl. After 3, 7, and 14 days, samples were fixed in formalin (Fisher Scientific, Fair Lawn, NJ) for 24 hours, then rinsed and stored in PBS until imaging. Cell-seeded scaffolds were permeabilized with 0.2% (v/v) Triton X-100 for 15 min and washed with PBS again. Cell nuclei and filamentous (F)-actin were labeled with 6-Diamidino-2-phenyindole dilactate (DAPI) and Tetramethylrhodamine Isothiocyanate (TRITC) at 1:1000 and 1:300 dilutions in PBS for 15 minutes and 1 hour, respectively. Labeled cells fixed to 3D scaffolds were observed via fluorescence (Axio Zoom V16 Fluorescence Microscope, Zeiss, Oberkochen, Germany) and confocal (Andor W1 Spinning Disc Confocal Microscope, Oxford Instruments, Abingdam, United Kingdom) microscopy. Z-stack images were processed and analyzed with ImageJ (NIH).

#### 2.5.4 Cell infiltration

Cryogels, previously stored in PBS, were submerged in individual 5 mL Eppendorf tubes containing a 30% w/v sucrose cryoprotectant solution and refrigerated at 4°C for 24 hours. After cryoprotection, a 5% w/w gelatin and 5% w/w sucrose embedding solution was prepared and dissolved in a water bath. Embedding molds were filled with 1 mL of the embedding solution and cryoprotected samples were transferred to their respective molds with additional solution added for complete coverage. The molds were incubated at 45°C for 2 hours, then frozen overnight at −80°C. Samples were then cryosectioned at 40-50 μm thickness (Cryostat Microm HM 525, Thermo Scientific, Waltham, MA, USA).

#### 2.5.5 Histology

All fixed samples were processed using an automated paraffin tissue processor (Tissue-Tek VIP 5, Sakura, Torrance, California, USA) for 8 hours. Briefly, samples underwent sequential dehydration through graded alcohols (70%, 95%, and 100%), followed by three changes of xylene and a four-stage paraffin infiltration step, and then embedded in paraffin blocks for sectioning. The embedded samples were sectioned at 4 µm using a microtome, floated in 47°C water bath to ensure smoothness, transferred to adhesive slides, and subsequently air-dried. All slides were then heated at 60°C for a minimum of 30 minutes to remove residual moisture. Paraffin was removed via two, five-minute immersions in xylene, followed by rehydration through graded alcohols (100%, 95%) into distilled water. The slides were then immersed in a silver nitrate solution and exposed to bright sunlight for 10–20 minutes, where the reaction was monitored periodically and terminated once calcium deposits turned brown/black. Following this, the sections were rinsed thoroughly in distilled water and treated with 5% sodium thiosulfate for 2–3 minutes. After another rinse in distilled water, the sections were counterstained with nuclear-fast red for 5 minutes. Finally, the slides were washed in distilled water, dehydrated through graded alcohols, cleared with xylene, and mounted with coverslips for microscopic examination.

#### 2.5.6 Critical Point Dryer

All cellularized and mineralized scaffolds were processed by critical point drying (SAMDRI®-795, Tousimis, Rockville, MD) in preparation for SEM imaging and histology.

#### 2.5.7 MicroCT

A hydroxyapatite lattice was imaged using the Bruker SkyScan 1172 micro-CT system (Billerica, MA, USA) at 68 kV and 145 µA, equipped with a 0.5 mm aluminum filter. The microCT datasets were reconstructed with NRecon software, employing a dynamic range of 0.1, a ring artifact correction of 10, and a beam hardening correction of 20%, resulting in a pixel size of 9.86 µm.

### 2.6 Statistical Analysis

GraphPad Prism was utilized to conduct all statistical analyses, with a significance level of 0.05. A two-way ANOVA, followed by a Tukey post hoc analysis, was performed to quantify the significance between groups from compression testing. Outliers in the swelling ratio were determined using the ROUT method, with a Q value of 1%.

## 3. Results and Discussion

### 3.1 Characterization of thermal treated 3D-printed lattices

Using a previously developed procedure, a gyroid lattice 3D model was created from MSLattice and Blender (Blender Institute, Amsterdam, Netherlands) with the following dimensions: 10 mm height, 9 mm diameter, 20% relative density, and 2.5 mm unit cell size.^22^ Note that the gyroid structure with these dimensions was chosen due to its desirable swelling ratio and mechanical properties when integrated with a cryogel scaffold. The gyroid was printed on a DLP 3D-printer (Fig. 1A) using HA-WOL ceramic resin. The gyroid structure (Fig. 1B, left) was selected because of its interconnected porous structure and high surface area-to-volume ratio, which supports cell attachment, proliferation, and differentiation, as well as angiogenesis, making it ideal for tissue engineering.^46,47^ Following printing and UV light curing, SEM images of the green lattices revealed distinct topographical features of spherical HA particles interspersed with lamellar and fibrous WOL particles (Fig 1B, right). Initially probed with TGA to determine the temperature required to combust most of the binder in the structure (Supplemental Fig. 1), lattices were then treated with two different thermal processes: Cycle 1 (C1) or Cycle 2 (C2). SEM images revealed that the ceramic HA and WOL particles appear to decrease in density for C1 lattices, while retaining the distinct morphology seen in green samples. This suggests that the ceramic particles are loosening and falling away due to weaker bonding between them, supporting that thermal treatment is burning the binder material set to hold the print together. Comparatively, C2 lattices display complete morphological changes in their surface, with what appears to be both HA and WOL particles fused together, indicating that the lattice achieved temperatures promoting particle sintering and conversion to intermediate mineral species (Fig. 1C).

**Fig. 1.**
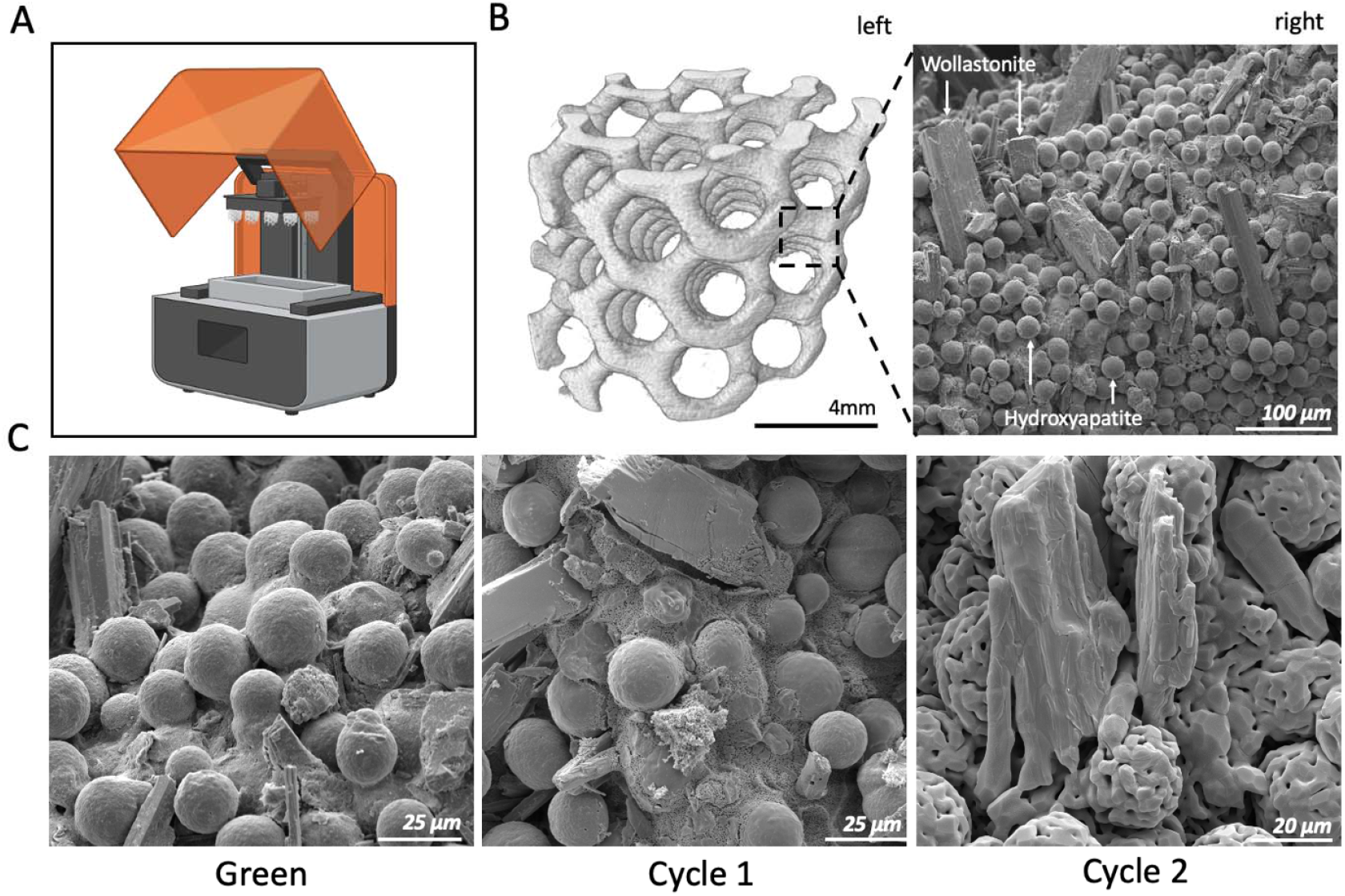
Fabrication and thermal treatment of ceramic gyroid lattices. A) DLP 3D-printed resin-based ceramic lattices (Biorender); B) Left: 3D reconstructed microCT of green ceramic lattice. Right: surface SEM image of green ceramic lattice; C) SEM images of green (Grn), Cycle 1 (C1), and Cycle 2 (C2) treated lattices.

FTIR further indicates the loss of organic material due to decomposition of the resin binder during the thermal treatment. The absorption peak corresponding to the C=O (∼1735 cm^-1^) bond in common acrylate and polyurethane polymer resins is no longer present after the C1 and C2 processes, with no other peaks indicating the presence of organic material, confirming successful decomposition of the binder (Fig. 2A, left). Additionally, FTIR spectra of C2 lattices show absorbance peaks at 1010 and 1050 cm□^1^, characteristic of both phosphate and silicate groups, as well as peaks at 880 and 930 cm□^1^, which are indicative of stretching vibrations of Si–O bonds in silicate groups (Fig. 2A, right).^48–50^ These characteristic peaks are consistent with the superimposition of the individual spectra of WOL and HA, further confirming the presence of both components within the resin, as their distinct molecular vibrations contribute to the observed spectral profile.^51,52^ X-ray diffraction illustrates the morphological changes of the mineral structure left behind following decomposition of the organic binder. Green and C1 lattices show peaks confirming the presence of hexagonal HA and triclinic WOL crystal structures (Fig. 2B). C2 lattices exhibit additional peaks corresponding to β-tricalcium phosphate (β-TCP) and silicocarnotite crystal structures, indicating the decomposition of HA and WOL into additional calcium phosphosilicate species. Debye-Scherrer analysis of mineral peaks yields spherical grain size estimates of 10-50 nm, on the same order of magnitude as mineral crystallite size in human bone, which may promote cell proliferation more effectively than micrometer-scale mineral grains (Equation 1).^53–56^ Compression tests (Table 1) confirm successful sintering, with C2 lattices achieving compressive moduli values roughly three times greater than green structures. C2 lattices additionally demonstrated significant increases in mechanical integrity when compared to C1 lattices, with a compressive modulus of 15.21 ± 1.83 MPa and an ultimate strength of 113.24 ± 10.31 kPa (Fig. 2C and Table 1). Bulk material compression tests revealed similar results, where C1 treated material achieved a 53.51 ± 8.37 MPa compressive modulus and 1.54 ± 0.37 MPa ultimate strength, whereas C2 treated materials achieved a 127.66 ± 61.50 MPa compressive modulus and 4.28 ± 1.56 MPa ultimate strength (Fig. 2D and Table 1). All three samples for C1 and two samples from C2 exhibited a plateau region at 1 MPa; this suggests a reorganization of the microstructure of the bulk samples, likely due to collapsing of pores. A plateau region is absent from the third sample of C2 which may also explain the deviation of its compressive modulus from the other two samples, causing a high standard deviation for compressive modulus. Burnout variability, sintering variability, or particle distribution may explain the differences with the third sample of C2.

**Table 1.** Compressive properties of green, thermally treated (C1 and C2) lattices, and bulk material cylinders.

|  | Lattice |  |  | Bulk Material |  |  |
| --- | --- | --- | --- | --- | --- | --- |
|  | Green | Cycle 1 | Cycle 2 | Green | Cycle 1 | Cycle 2 |
| Compressive Modulus | $5.38 \pm 1.27$ MPa | $1.62 \pm 0.60$ MPa | $15.21 \pm 1.83$ MPa | $668.00 \pm 79.19$ MPa | $53.51 \pm 8.37$ MPa | $127.66 \pm 61.50$ MPa |
| Linear Strain Range | $0.0269 \pm 0.0103$ | $0.0060 \pm 0.0019$ | $0.0075 \pm 0.0013$ | $0.0256 \pm 0.0033$ | $0.0214 \pm 0.0085$ | $0.0096 \pm 0.0052$ |
| Ultimate Strength | $170.00 \pm 4.03$ kPa | $8.40 \pm 0.44$ kPa | $113.24 \pm 10.31$ kPa | $66.49 \pm 2.49$ MPa | $1.54 \pm 0.37$ MPa | $4.28 \pm 1.56$ MPa |
| Ultimate Strain | $0.0572 \pm 0.0031$ | $0.0069 \pm 0.0033$ | $0.0412 \pm 0.0086$ | $0.2247 \pm 0.0230$ | $0.0666 \pm 0.0278$ | $0.0539 \pm 0.0167$ |

**Fig. 2.**
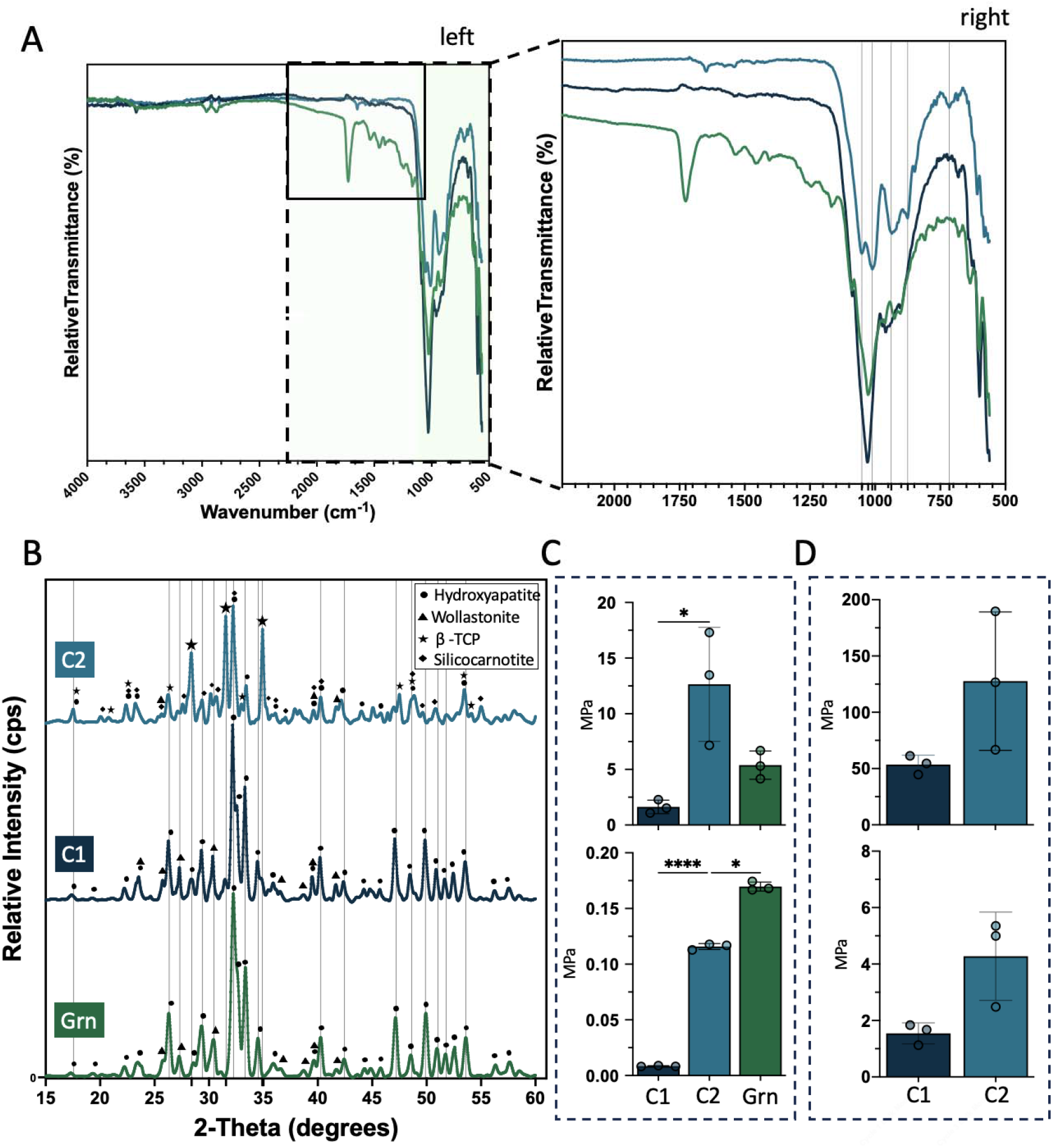
Physical and chemical characterization of thermal treated 3D-printed ceramic lattices. A) Left: FTIR spectra of green, Cycle 1 (C1), and Cycle 2 (C2) lattices. Right: Relative transmittance spectra; B) XRD spectra profiles and peaks matched using Jade software; Compressive modulus (top) and ultimate strength (bottom) values of C) lattices and D) bulk material.

### 3.2 Biocompatibility and cytotoxicity characterization of green and sintered lattices

To assess cell viability within 3D-printed lattices, green and C2 sintered lattices were cell seeded and imaged for biocompatibility, where C2 was selected for its superior mechanical characteristics and crystalline composition. All lattices were cultured in non-treated tissue culture, plates (NT-TCP), whereas treated plates (T-TCP) were seeded and used as controls. Following 6, 24, and 48 hours of culture and subsequent Live/Dead staining, live cells were adhered to sintered lattices at all time points, whereas no live cells were visibly adhered to green lattices (Fig. 3A). When imaging lattices left within TCP wells, live cells were seen both migrating and beginning to adhere to the sintered lattice, which is slightly fluorescent in the red wavelength. Only a few dead cells are visible in both control and sintered TCP wells, indicated by the positive stain (red) in the image (Fig. 3B). In comparison, the cells seeded on green lattices are shown migrating from the lattice, with what visually appears to be more than 50% cell death at 24 hours, and almost no visually living cells in the TCP well at 48 hours (Fig. 3B). The CCK-8 assay further confirms these results, with evidence of reduced cell activity with the green lattices when compared to sintered lattices and a control at every time point (Fig. 3C top). Moreover, when normalized to a control, cell viability percentage decreased overtime and was found to be significantly reduced in green lattices when compared to sintered lattices, which maintained viability around 50% from 24 to 48 hours, indicating cell proliferation rates matching that of a control (Fig. 3C bottom). Finally, after 72 hours of culture, lattices fixed and stained to visualize cell nuclei and f-actin were imaged, depicting cellular activity and attachment with blue nuclei and orange f-actin throughout the sintered lattice (Fig. 3D). Note that, while there appears to be no distinct cellular features on the green lattice, the structure is auto-fluorescent in both channels. However, these results strongly suggest that the green lattice is cytotoxic at early time points, while the sintered lattice is biocompatible, supporting cellular proliferation and attachment up to 72 hours. These results demonstrate that sintering DLP 3D-printed ceramic structures enables biocompatibility and eliminates cell death from resin.

**Fig. 3.**
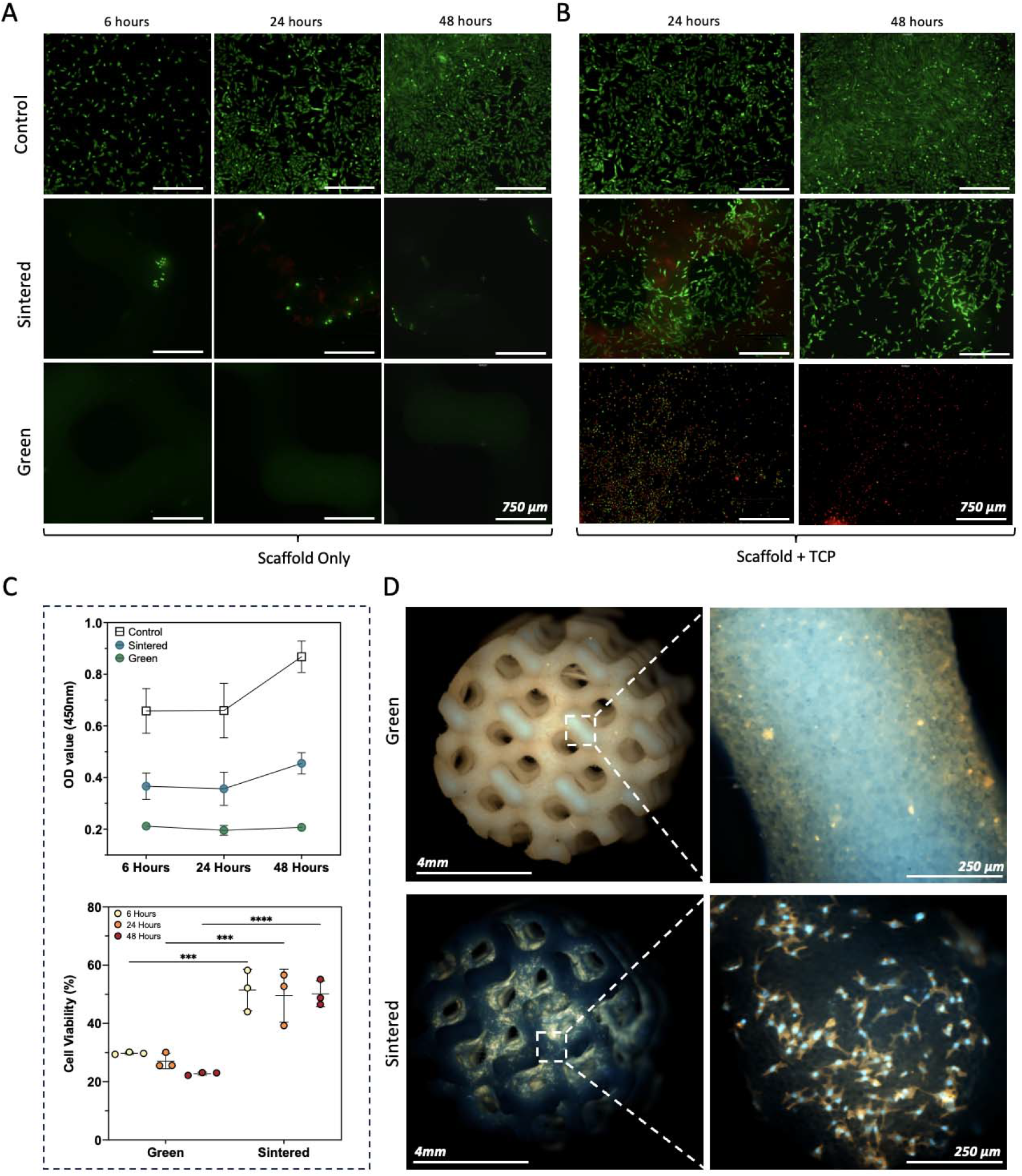
Early biocompatibility and cytotoxicity analysis of 3D-printed ceramic lattices. A) Live/Dead stain depicting cell viability (live cells appear green, dead cells appear red) of control well, green, and sintered 3D-printed lattice; B) 3D-printed lattice with TCP; C) CCK8 assay highlighting cell activity/viability or death of cells during co-culture with lattices; D) Fluorescence microscopy images of green and sintered lattices with cell nuclei (blue) and cytoskeleton (red/orange) visible after 72 hours of culture.

### 3.3 Sintering 3D-printed lattices enables biocompatibility in-vitro

Long term cell viability was tested by culturing MG63 cells on sintered or green lattices for 3,7, and 14 days. High resolution fluorescent confocal microscopy images of green lattices appear to depict no signs of cellular activity on the structures at any time point. Further, as green lattices fluoresce brightly at 405 nm wavelengths, actin structures were stained with tetramethylrhodamine (TRITC) and imaged at 561 nm. While there appears to be some red auto-fluorescence from the structure, there does not appear to be any adhered cellular structures, with SEM images confirming these results with no visible cells on its surface after 3,7, and 14 days of co-culture (Fig. 4A, top row). In comparison, confocal and SEM imaging of sintered lattices depict significant levels of cell proliferation and spreading, with adhered cells covering most of the lattice surface after 14 days of incubation (Fig. 4A, bottom row). Additionally, morphological features of single cells depict strong cellular attachment and tension with multiple focal adhesion points, wide spreading, and visible actin stress fibers, indicating that the cells are healthy and interacting strongly with the material at all time points (Fig. 4B). Furthermore, there is evidence of advanced gross cellular organization, with clumping and polarized morphology along and across distinct lattice features, suggesting that the sintered material and lattice structure may be prompting some directed cellular migration (Supplemental Fig. 2 and Video 1). This could be explained by cell morphological features visible in the SEM images that show evidence of contact guidance, with cells aligning and anchoring within grooves and taking topographical cues from the 3D-printed lattice (Supplemental Fig. 3). These findings further confirm that sintering DLP 3D-printed ceramic structures enables biocompatibility, with evidence that it is also bioactive.

**Fig. 4.**
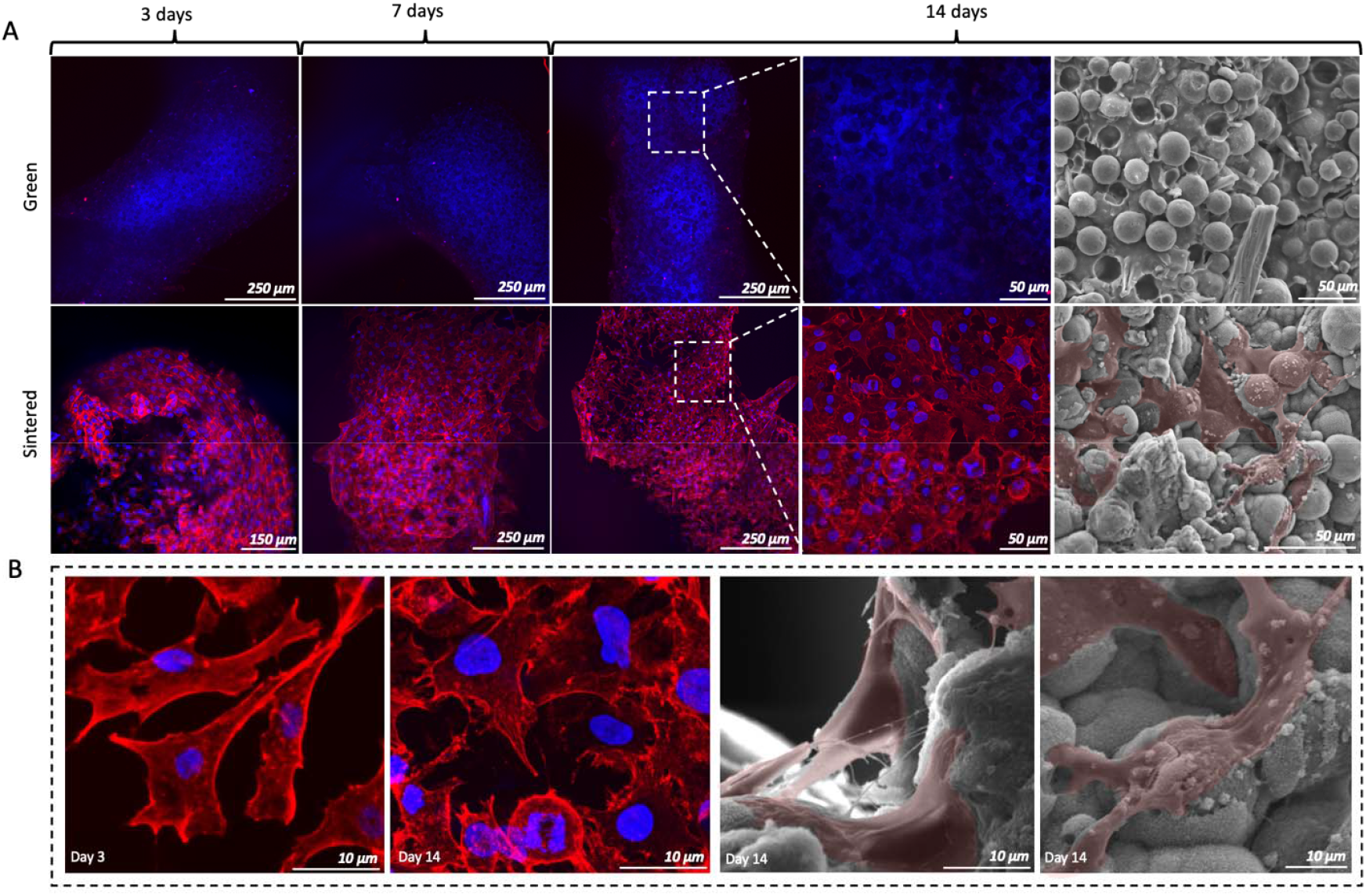
Long term cellular viability and cell morphology on sintered 3D-printed ceramic lattices. A) Confocal and SEM images* of green (top row) and sintered lattices (bottom row) co-cultured with MG63 cells and stained with DAPI and TRITC to visualize nuclei (blue) and actin filaments (red); B) Higher magnification images comparing cellular structures on sintered lattices on day 3 and 14 of incubation. *Cells were manually highlighted red for visualization.

### 3.4 The addition of sintered lattices significantly enhances polymer cryogel mechanical properties

SEM imaging was used to capture scaffold microstructures and PoreVision was used to analyze pore areas (Figure 5A).^44^ No significant differences in pore areas were observed between the cryogel and composite scaffolds (p>0.05). While both scaffolds exhibited a skew towards smaller pores, larger pore formation was also noted. Smaller pores predominated due to a high nucleation density and freezing conditions; however, regions with fewer nucleation sites, slower freezing rates, or lower polymer concentration allowed individual ice crystals to grow, resulting in larger pores. Despite the skew towards smaller pores, larger pores also formed in regions with fewer nucleation sites and other localized variations, resulting in a distribution of both small and large pores. SEM images (Supplemental Fig. 4) revealed that incorporating cryogel into a 3D-printed ceramic lattice altered its pore architecture, with attachment points between the cryogel and lattice indicating interaction between the components. Notably, pores farther from the lattice appeared larger, while those near the attachment points were smaller. This pattern likely resulted from the ceramic lattice hindering ice crystal growth and creating nucleation sites that facilitated the formation of smaller ice crystals near the lattice, while fewer nucleation sites farther from the lattice allowed larger ice crystals to form. The pores exhibited a visually ovoid shape, and the isoperimetric ratio of 0.42 quantitatively supports this, suggesting an elongated, jagged structure like the trabecular bone architecture observed in previous studies.^57–59^ Despite these variations, the addition of lattice did not significantly affect the average size in the composite scaffolds. Notably, maintaining the porosity of the cryogel is also crucial for successful incorporation into bone defect sites, as pore size and interconnectivity are essential for cell attachment, nutrient transport, and growth factor delivery. Previous research indicates that a pore diameter of 100– 200 μm is optimal for bone regeneration.^60,61^ Although the mean pore size of composite scaffolds in this study fell below the optimal range, they were comparable to plain cryogel scaffolds (Fig. 5A). Therefore, we hypothesize that pore shrinkage occurred during lyophilization and it is likely that the pores in the hydrated scaffolds are actually larger due to successful cell seeding and viability. Future research should investigate the pore size in its hydrated state using microCT.

**Fig. 5.**
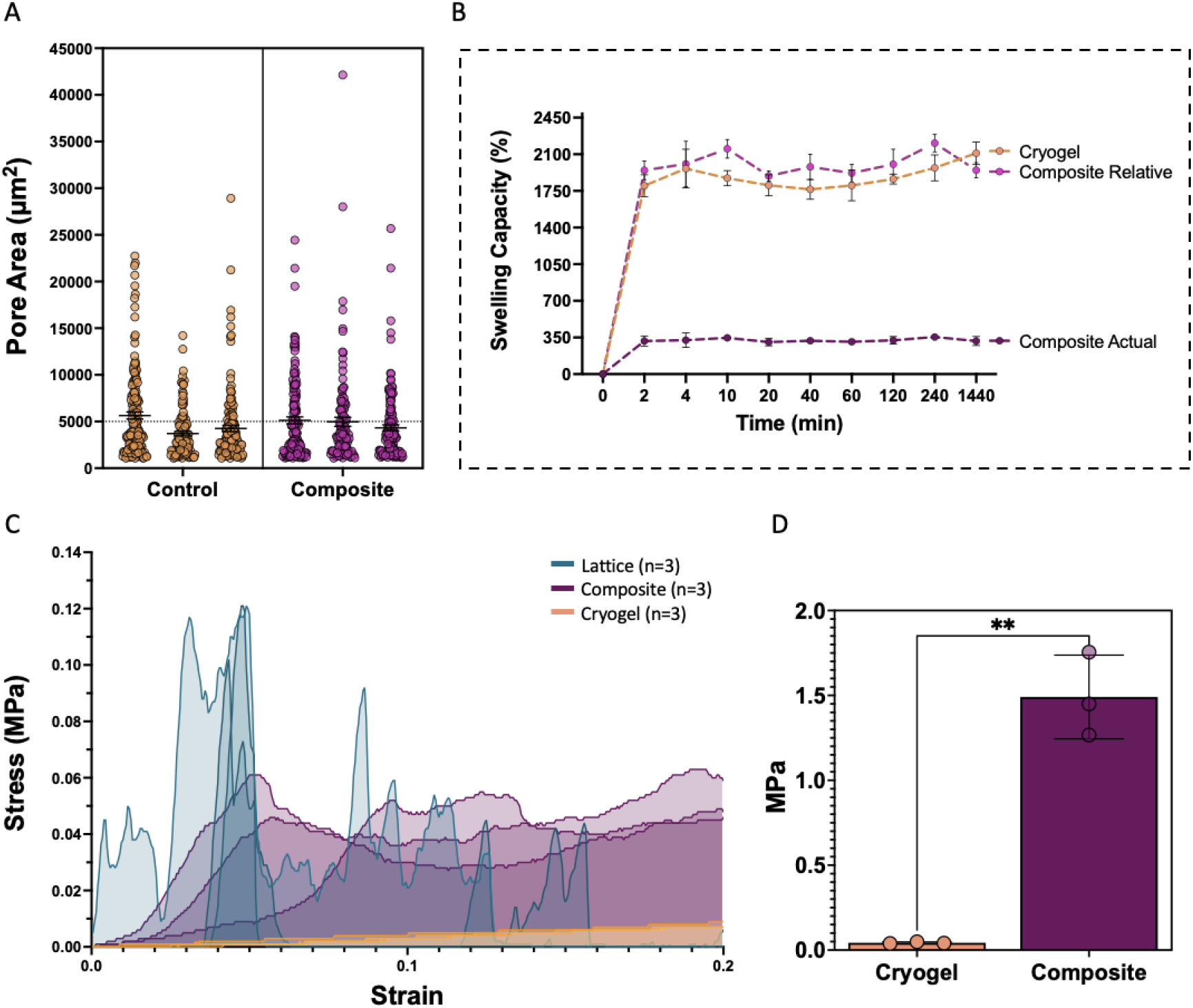
Physical and mechanical properties of composite and cryogel scaffolds. A) Pore area distribution within both cryogel and composite scaffolds; B) Swelling capacity of cryogel (cryogel), composite scaffolds (composite actual), and composite scaffolds relative to the amount of cryogel within them (composite relative); C) Stress–strain curves for lattices (blue), composite scaffolds (purple), and cryogel scaffolds (orange); D) Compressive modulus for cryogel and composite scaffolds.

Following dehydration and subsequent immersion in water, the cryogel and composite scaffolds reached their relative maximum swelling capacity within 2 minutes, maintaining approximately 1950% swell capacity for up to 24 hours. To accurately determine the swelling ratio of the cryogel within the composite scaffold, the weight of the ceramic lattice was subtracted from the total scaffold weight (Equation 3), ensuring that only the cryogel’s (relative) contribution was measured (Fig. 5B). No significant differences in swelling ratio were observed within the cryogel timepoints between the 2-minute mark and later, within the composite timepoints between the 2-minute mark and later, or within the composite relative timepoints between the 2-minute mark and later (Q = 1%). There was no statistical difference between the cryogel and composite’s relative swelling capacity. Whereas the swelling ratio of the composite group was statistically different at all timepoints compared to both the cryogel and composite relative groups (p<0.05). Rapid swelling is a beneficial property for scaffolds in bone tissue engineering as it facilitates nutrient absorption and promotes uniform cell distribution, enhancing tissue regeneration. In this study, both cryogel and composite scaffolds achieved maximum swelling capacity within two minutes, where the addition of the 3D-printed ceramic lattice did not reduce the relative swelling capacity compared to cryogel scaffolds.

Scaffolds were compressed to 50% strain to evaluate their mechanical properties (Figure 5C, Table 2). The stress–strain curves for the cryogel scaffolds exhibited a continuous increase in slope without ultimate failure, a characteristic behavior of polymer compression. In contrast, the composite scaffolds displayed multiple peaks in their stress–strain curves, corresponding to the sequential failure of lattice layers under high loads (Figure 5C). The composite scaffolds exhibited a significantly higher average compressive modulus (1.49 ± 0.24 MPa) compared to the cryogel scaffolds (0.04 ± 0.01 MPa; Figure 5D; p<0.05). Additionally, the cryogel scaffolds exhibited a broader linear strain range (21.70 ± 4.12%) compared to the composite scaffolds (2.25 ± 0.47%), highlighting their exceptional elasticity and ability to deform without failure. Cryogels, known for their highly porous, sponge-like structure and remarkable resilience, can withstand compression up to 90% strain and fully rebound without crack propagation.^13,14,16^ In contrast, the composite scaffolds, with their integrated ceramic lattice, achieved an ultimate strength of 53.30 ± 7.42 kPa and an ultimate strain of 6.82 ± 2.39%, demonstrating their significantly enhanced load-bearing capacity and structural integrity. The cryogels endured a stress of 15.17 ± 0.47 kPa at 50% strain; since they did not fail at this strain, this cannot be referred to as the ultimate stress. These mechanical differences underscore the adaptability of cryogels for applications requiring high elasticity and flexibility, while the composite scaffolds are better suited for bone defect sites that may experience greater relative loading. The improved mechanical properties of the composite scaffolds, coupled with their controlled failure mechanisms, make them a promising option for bridging the gap between stiffness and adaptability in bone tissue engineering applications.

**Table 2.** Compressive properties of cryogel and composite scaffolds (n = 3).

|  | **Cryogel** | **Composite** |
| --- | --- | --- |
| **Compressive Modulus (MPa)** | 0.04 ± 0.01 | 1.49 ± 0.24 |
| **Linear Strain Range** | 21.70 ± 4.12% | 2.25 ± 0.47% |
| **Ultimate Strength (kPa)** | N/A | 53.30 ± 7.42 |
| **Ultimate Strain** | N/A | 6.82 ± 2.39% |

### 3.5 Composite scaffolds exhibit enhanced bioactivity

To compare cellular interactions and activity on cryogel and composite scaffolds, MG63 (osteoblast-like) cells were seeded and cultured on samples for 3, 7, and 14 days. Confocal imaging shows cellular activity on the surfaces of both cryogel and composite samples at all time points, with the highest number of cells at the 7-day timepoint for both samples, where composite scaffolds (µ=1321 cells) have roughly 5 times the number of cells on their surface compared to cryogel samples (µ=295 cells) (Fig. 6A,B and Supplemental Fig. 5). Additionally, sectioned samples show cellular infiltration into the center of all samples at all time points, indicating cells are successfully proliferating throughout the scaffolds (Fig. 6C).

**Fig. 6.**
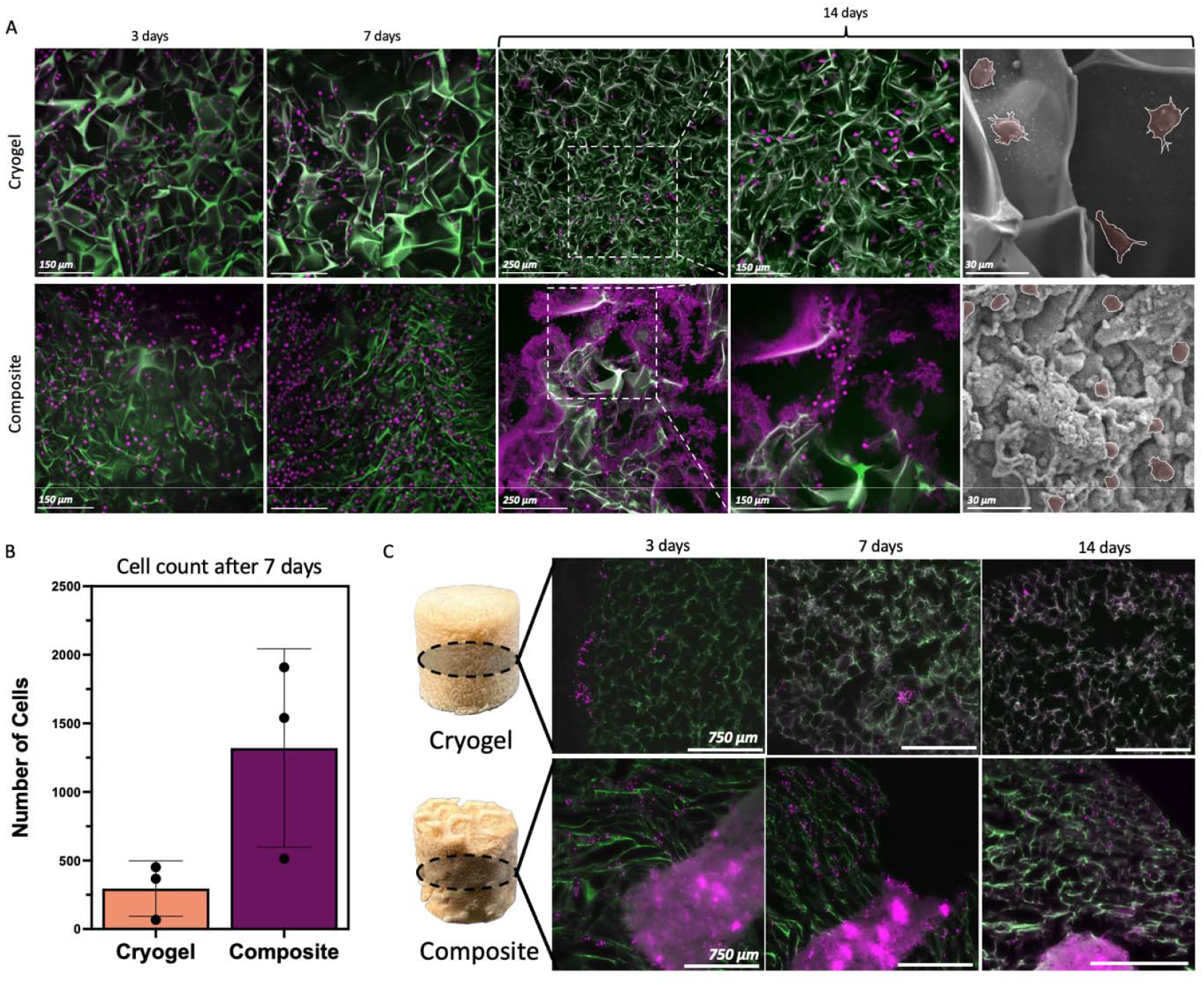
Long-term cellular interactions with cryogel and composite scaffolds. A) Confocal and SEM images of cryogel (top row) and composite (bottom row) scaffolds at 3-,7-, and 14-day timepoints of cellular incubation; B) Quantitative analysis of cells at 7 days (n=3)*; C) Fluorescent microscope images of sample cross-sections at 3-, 7-, and 14-days of cellular incubation. Note that cryogel appears green, while cell nuclei and mineral growth appear magenta. *Standard deviation error bar (p>0.05).

Interestingly, there is an apparent reduction in cell numbers at 14 days for both groups. With evidence of distinct fluorescent discoloration and material breakdown in the images, the drop in cell count is likely due to the degradation of the cryogel matrix, resulting in structural changes and mechanically instable conditions that lead to loss of surface integrity and collapses in the material matrix (Supplemental Fig. 6). Glutaraldehyde was used as the crosslinker for all cryogel scaffolds. Over time, glutaraldehyde crosslinking have been shown to become unstable in aqueous conditions and biological solutions, which would result in structural breakdown of polymer chains, leading to changes in the material that could alter the fluorescence intensity and distribution of the cryogel material.^62^ Additionally, late-stage cell reduction could also be a result of degradation byproducts from the breakdown of the glutaraldehyde, which can disrupt cellular processes and induce oxidative stress.^62–64^ Ongoing work will investigate the degradation and byproduct profiles of the formulated cryogels in this study to improve long term stability and biocompatibility of the gel.

Additionally, some mineral growth can be noted throughout the cell-seeded composite scaffolds, visibly auto fluorescing magenta under 405nm wavelength light in the confocal and cross-section images; this was also seen depicting distinct crystalline features in SEM images on day 14 (Fig. 6A, Supplemental Fig. 6). This mineral growth is thought to be occurring throughout the composite scaffolds via a combination of processes involving both cell- and surface reactivity-mediated mineralization. To investigate these processes further, cryogel and composite scaffolds underwent various treatments to test mineral induction and growth.

SEM analysis of cryogel samples revealed minimal mineral growth in both samples incubated in media for 14 days (media-incubated) and mineral-induction-soaked conditions, with sparse, small-scale mineral deposits primarily confined to the periphery, as indicated by faint positive Von Kossa staining (Fig. 7A). This suggests that while calcium-phosphate nucleation is occurring on a small scale, it is confined to the outer surfaces, likely due to limited ion penetration into the cryogel interior. In contrast, composite scaffolds exhibited distinct differences in mineralization patterns. Media-incubated composite scaffolds showed some mineral nucleation with small crystal clumps and thin peripheral mineral deposits, evidenced by light Von Kossa staining. Conversely, mineral-induction-soaked composite scaffolds displayed substantial mineralization, characterized by larger, uniform crystalline sheets and blocks with dark, thick Von Kossa-stained regions along the periphery, suggesting that the ceramic lattice provides an ion rich environment for mineral precipitation. However, without any extremal stimuli, ion concentrations, and therefore mineral growth, are still limited to the edges of the sample. On the other hand, composite scaffolds cell-seeded and -incubated for 14 days (cell & media incubated) exhibited high mineralization levels that depict a different morphology. SEM images reveal amorphous, dune-like rounded crystalline structures with clumped growths varying in size and form (Fig 6A. bottom row). This observed growth behavior and morphology of apatite agglomerates match those reported in previous work studying bioactive calcium-phosphate and calcium-silicate ceramics, including Silicocarnotite.^65–67^ Additionally, characteristic of biomineralization, these mineral structures may likely reflect cell-driven processes, where cells, proteins, and organic molecules disrupt the uninform crystalline structures apparent in mineral-induction-soaked samples.^68^ Histological analysis further corroborates these findings, with evidence of dark, positive Von Kossa stains throughout the entirety of the scaffold, suggesting scaffold-wide mineralization mediated by cell-driven processes (Fig. 7B, Supplemental Fig. 7). Quantitative Alizarin red stain measurements depict significant levels of calcium in all composite treated samples when compared with cryogel treated groups, while there were no significant differences in measured calcium between the treated composite samples (Fig. 7C). Bearing in mind that Alizarin stain binds nonspecifically to calcium, the lattice composition is made primarily of calcium and will therefore read higher when compared to cryogels. The similar Alizarin Red results suggest that total calcium content is comparable across composite samples, whereas the Von Kossa stain specifically identifies calcium bound in phosphate complexes, highlighting differences in more specific apatite formations across samples. These results depict the distinct roles of abiotic and cellular mechanisms in apatite formation and the bioactive influence of the ceramic lattice in supporting and enhancing both processes in the composite scaffold.

**Fig. 7.**
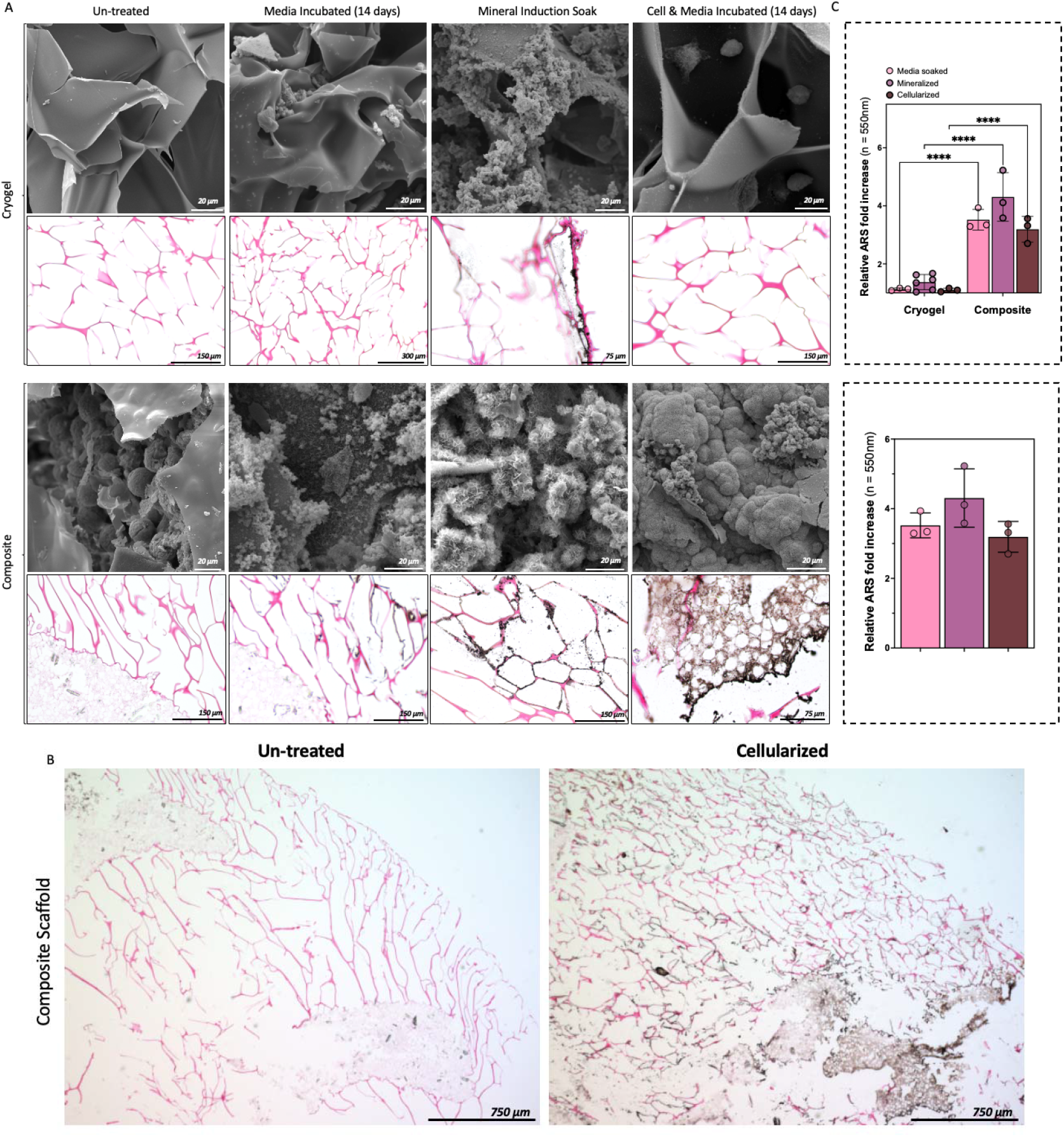
Enhanced mineralization potential of composite scaffolds. A) SEM and representative image of Von Kassa mineral stain of cryogel (top rows) and composite (bottom rows) left un-treated, incubated in cell media for 14 days, mineral induction soaked, and incubated with cells and media for 14 days; B) Sample-wide representative Von Kassa stain image of un-treated and cell incubated composite scaffolds; C) Quantitative analysis of Alizarin red stained samples.

## 4. Conclusion

This study demonstrates the successful fabrication of a composite scaffold for bone tissue engineering, integrating thermally treated DLP 3D-printed ceramic lattices with cryogel. The advanced gyroid lattice design, produced via DLP-based 3D-printing and thermal sintering, enabled precise control over pore architecture, enhanced mechanical properties, and improved biocompatibility. Thermal treatment cycles were shown to significantly influence the microstructure and material properties of the lattices, with Cycle 2 yielding sintered lattices that exhibited superior compressive modulus, ultimate strength, and phase transformation compared to bioactive β-TCP and Silicocarnotite. Evidenced through MG63 cell culture experiments, these structural and compositional features enabled effective cellular attachment, proliferation, and viability. This is a significant finding that demonstrates that the identified sintering cycle can transform more readily available and less expensive HA and WOL particles into bio-ceramic mixed composites that possess improved bioactivity and resorbability, creating constructs that are better suited for bone regeneration applications, all the while maintaining structural integrity. Then, when integrated with cryogel scaffolding, the resultant composite scaffolds retained their relative swelling capacities while exhibiting significantly enhanced mechanical performance and bioactive properties when compared to cryogel-only constructs. Mineral induction tests and *in vitro* studies depict strong evidence of these conclusions, with significant calcium-phosphate mineral deposits found throughout cellularized composite samples, and further evidence of cell proliferation and infiltration at all time points. However, since it appears that the cryogel is breaking down faster than anticipated in *in vitro* cell culture, future work will investigate and optimize the cryogel degradation mechanism and rate to improve its long-term compatibility with the lattice. Overall, the combination of bioactive ceramic lattice with cryogel bridges the gap between mechanical strength and elasticity, addressing critical challenges in bone defect repair. These composite scaffolds demonstrate promise for applications requiring high resolution patient specificity, structural integrity, and dynamic biological activity, making them a compelling platform for future studies aimed at enhancing bone regeneration. Further investigation into long-term composite stability, bioactive efficacy in inducing osteoprogenitors, and *in vivo* performance in complex bone defect repair will be pivotal in advancing this technology toward clinical applications.

## Data availability

The data that support this study are available within the article and its Supplementary data files, or available from the authors upon request.

## CRediT author contribution statement

**Peter M. Bertone:** Writing – Original Draft, Conceptualization, Methodology, Investigation, Visualization, Formal analysis, Funding acquisition. **Levi M. Olevsky:** Writing – Original Draft, Conceptualization, Methodology, Validation, Investigation. **Kavin Kathir:** Investigation, Writing – Review and Editing. **Simon A. Agnew:** Investigation, Writing – Review and Editing. **William J. Scheideler:** Conceptualization, Writing – Review and Editing. **Katherine R. Hixon:** Conceptualization, Methodology, Resources, Funding acquisition, Writing – Review and Editing.

## Acknowledgements

The authors would like to acknowledge Scott Palisoul, Dartmouth Biomedical Engineering Center, Life Sciences Light Microscopy facility at Dartmouth, Daniel Cullen, the Electron Microscope facility at Dartmouth, and Matthew Genser for their assistance with this manuscript.

