## Supplemental Figures for "Sintering 3D-Printed Hydroxyapatite-Wollastonite Lattices Improve Bioactivity and Mechanical Integrity for Bone Composite Scaffolds"

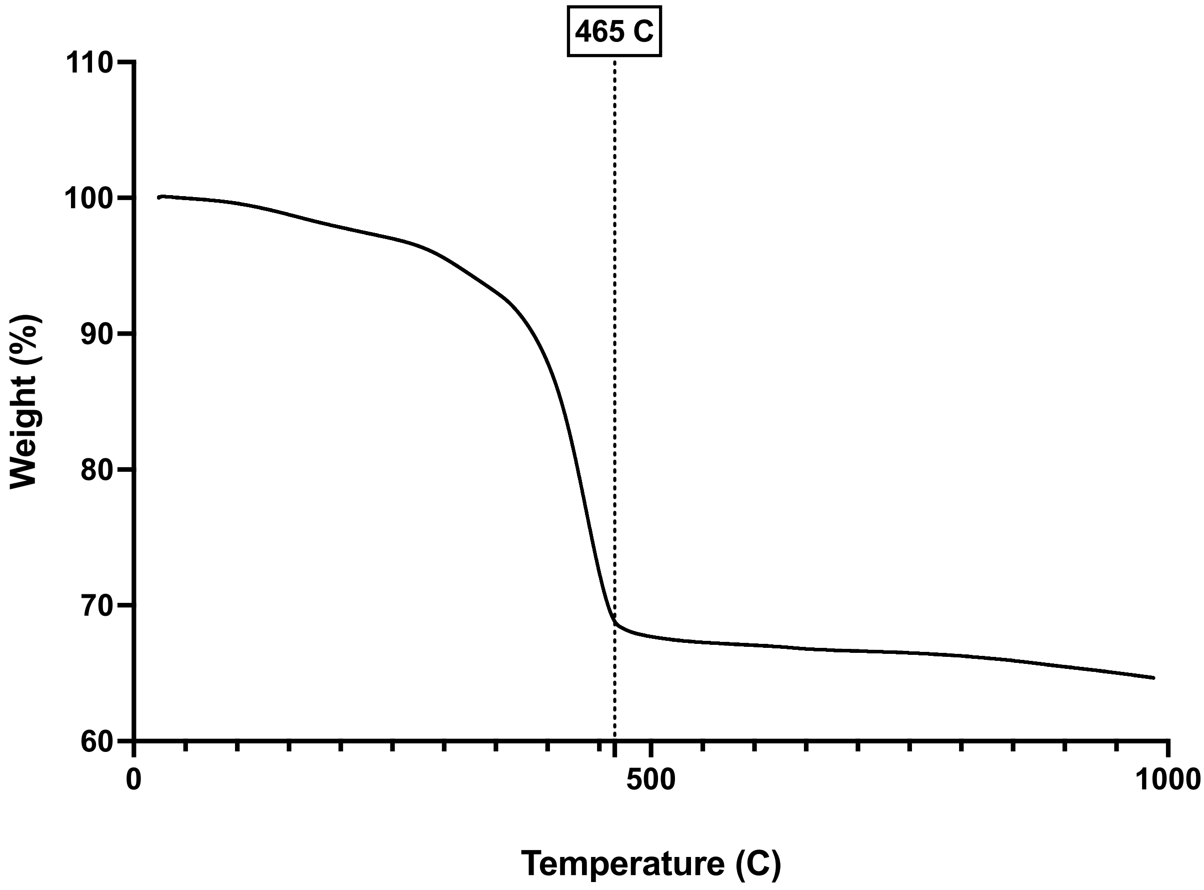


**Supplemental Figure 1**. Thermogravimetric analysis (TGA) measurement of 3D-printed HA-WOL lattice structure.

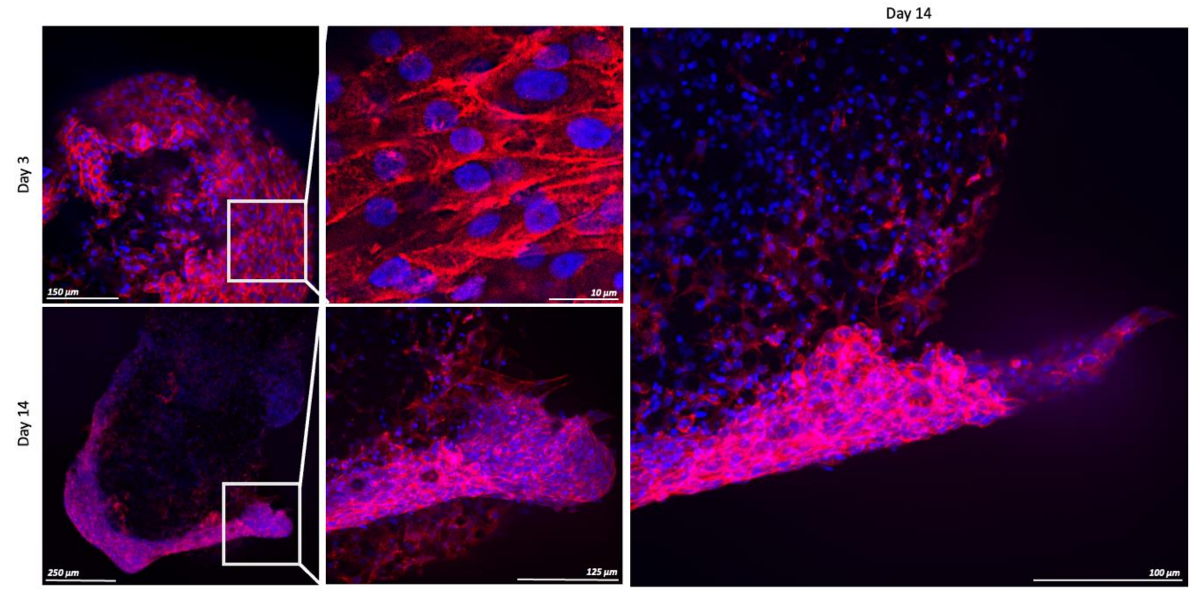


**Supplemental Figure 2.** Confocal microscopy images of MG63 cell growth on sintered lattices exhibiting aligned and clumping growth patterns. DAPI blue cell nuclei, TRITC Red cell actin.

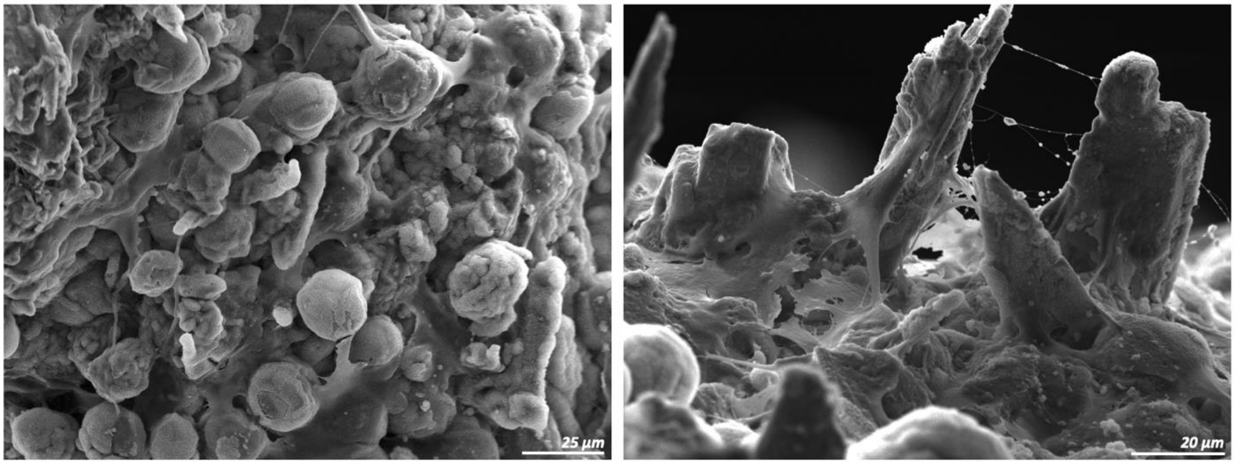


**Supplemental Figure 3.** SEM images of MG63 cell growth on sintered lattice surface after 14 days of culture.

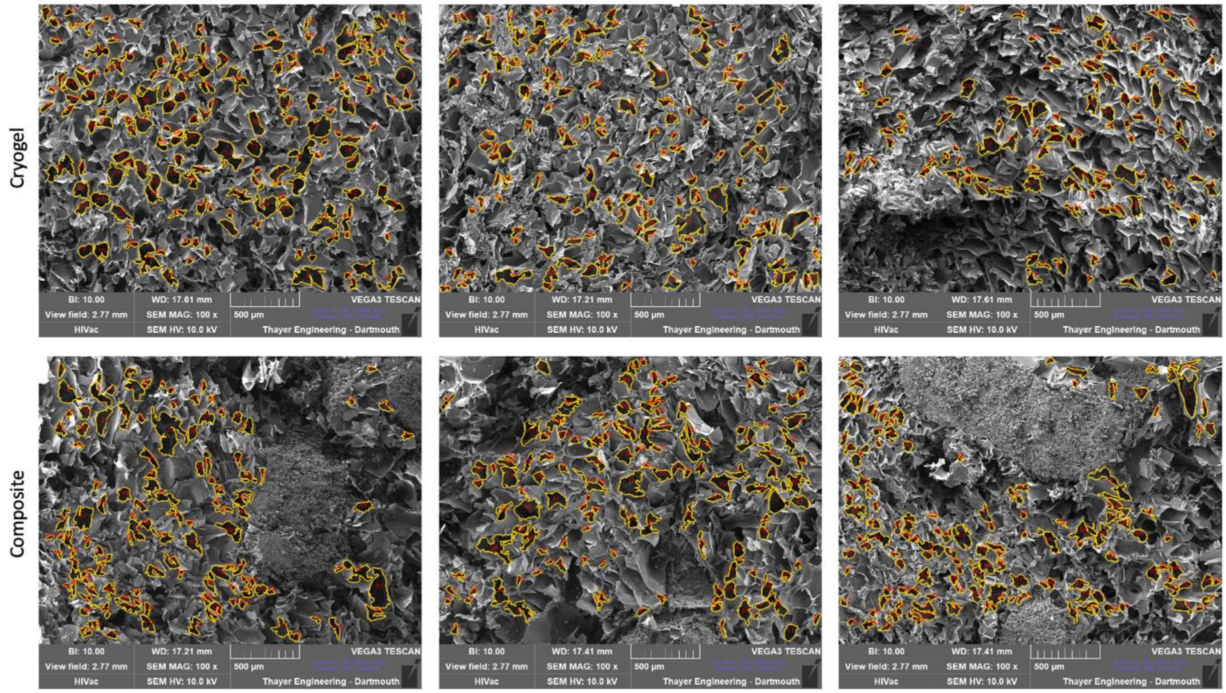


**Supplemental Figure 4.** SEM images of cryogel (n=3) and composite (n=3) samples overlayed with pore analysis software output isolating and measuring pore architecture (PoreVision).

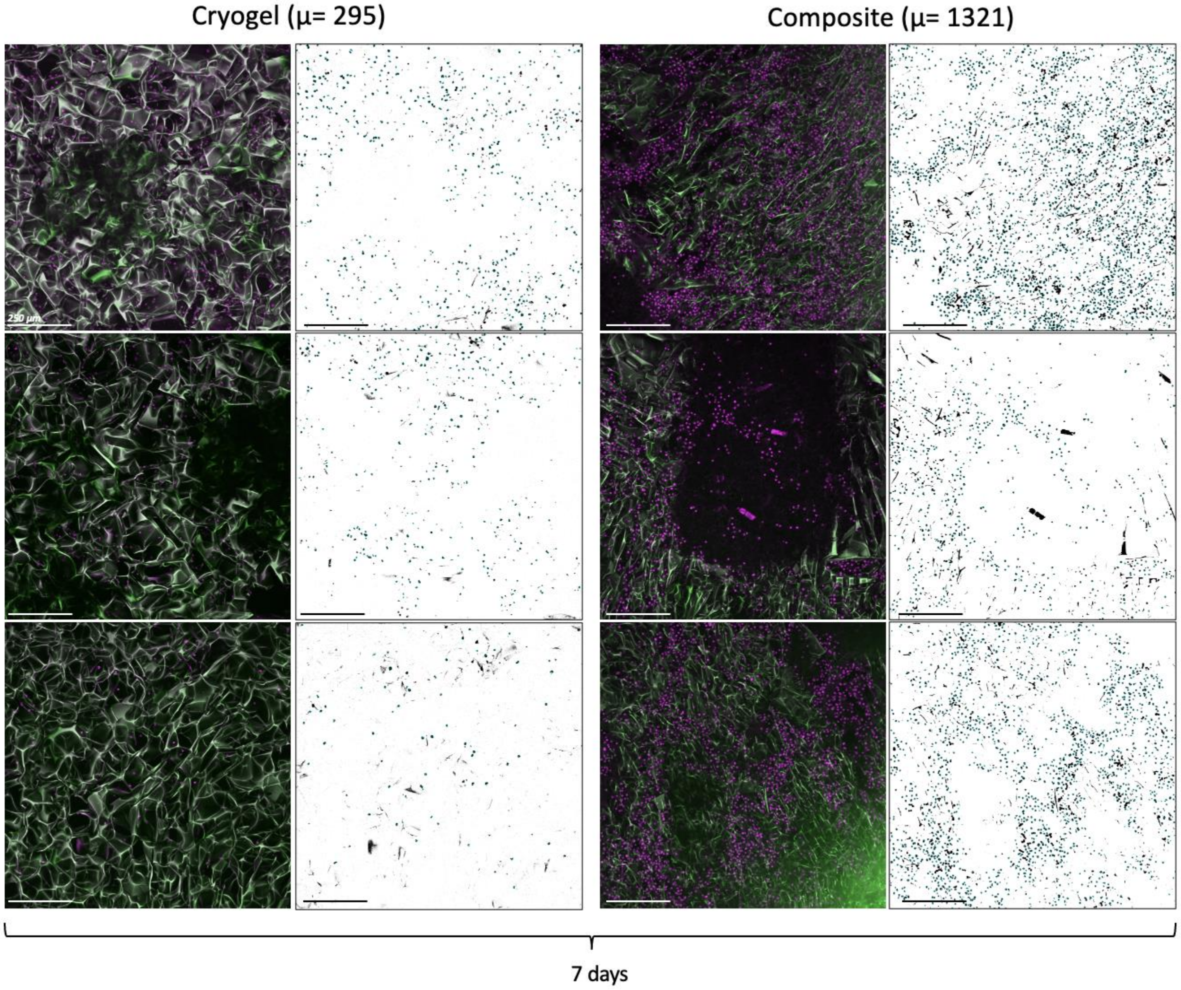


**Supplemental Figure 5.** Confocal images (left column) and ImageJ analysis isolating and counting individual cell nuclei (right column) for both cryogel (n=3) and composite (n=3) samples. Samples were imaged after 7 days of culture with MG63 cells stained with DAPI. Nuclei appear magenta, cryogel appears green.

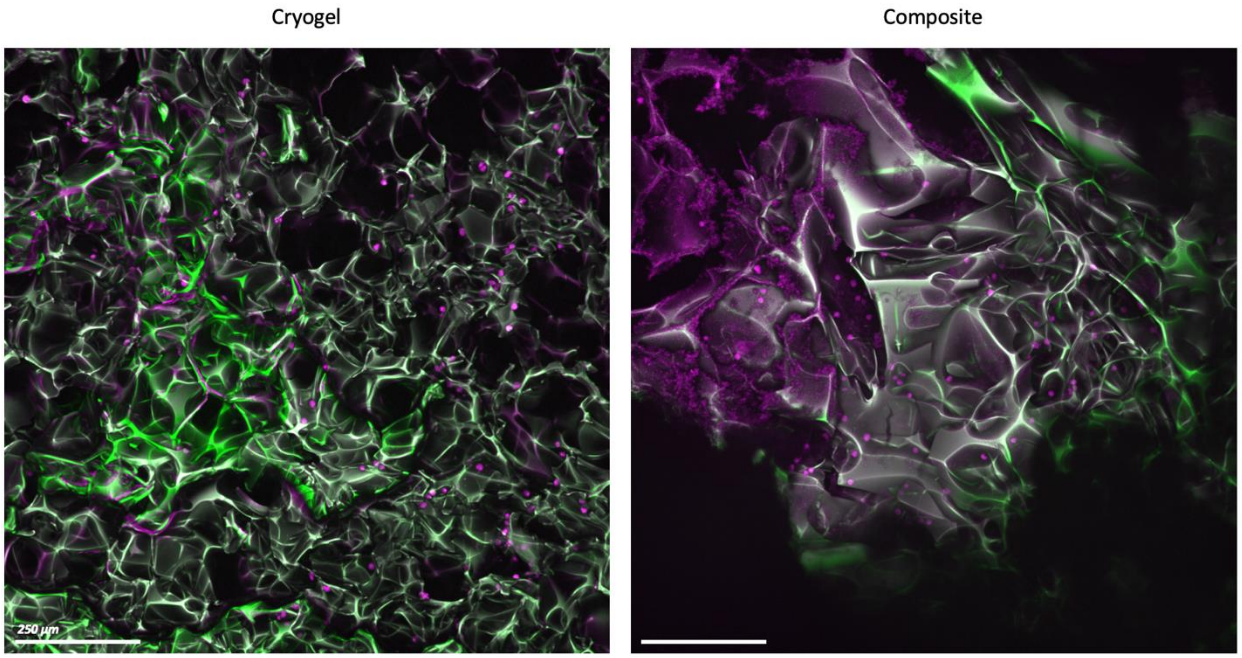


**Supplemental Figure 6.** Confocal images of cryogel (left) and composite (right) samples after 14 days of culture with MG63 cells. DAPI stained nuclei appear magenta, mineral growth appears magenta, and cryogel appears green.

Composite

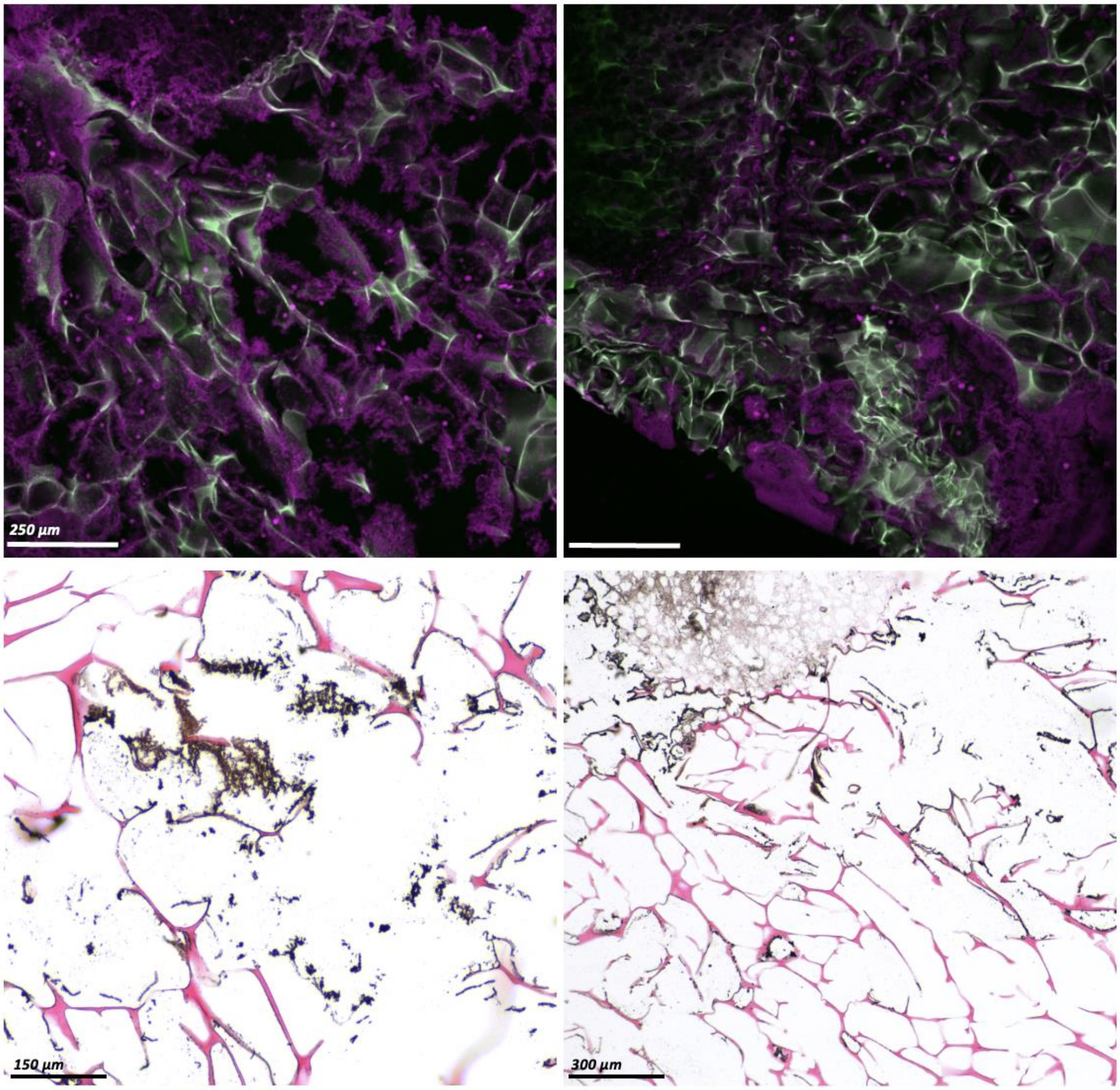


**Supplemental Figure 7.** Confocal images (top row) and histology slices stained with Von Kassa stain of composite samples after 14 days of culture and incubation with MG63 cells. DAPI stained nuclei appear magenta, mineral growth appears magenta, and cryogel appears green. For histology, mineral growth appears dark brown and black.
